# Active Microtubule-Actin Crosstalk Mediated by a Nesprin-2G-Kinesin Complex

**DOI:** 10.1101/2024.05.13.594030

**Authors:** Natalie Sahabandu, Kyoko Okada, Aisha Khan, Daniel Elnatan, Daniel A. Starr, Kassandra M. Ori-McKenney, G.W. Gant Luxton, Richard J. McKenney

## Abstract

Nesprins are integral membrane proteins that physically couple the nucleus and cytoskeleton. Nesprin-2 Giant (N2G) stands out for its extensive cytoplasmic domain, which contains tandem N-terminal actin-binding calponin-homology domains followed by >50 spectrin repeats and a C-terminal outer nuclear membrane-spanning KASH domain. N2G’s KASH domain interacts with the inner nuclear membrane, lamina-binding SUN proteins within the perinuclear space, forming a linker of nucleoskeleton and cytoskeleton (LINC) complex. Additionally, N2G contains a conserved W-acidic LEWD motif that enables the direct interaction with kinesin-1’s light chain, indicating N2G’s involvement with both actin and microtubules. The absence of N2G leads to embryonic lethality in mice, while cellular assays highlight N2G’s role in nuclear positioning across diverse biological contexts. However, the precise mechanisms underlying N2G-mediated nucleocytoskeletal coupling remain unclear. Here we study N2G’s interactions with F-actin and kinesin-1, revealing its functions as an F-actin bundler, a kinesin-1-activating adapter, and a mediator of active cytoskeletal crosstalk. Along with MAP7 proteins, N2G directly links active kinesin-1 motors to F-actin, facilitating actin transport along microtubule tracks. These findings shed light on N2G’s dynamic role as a crosslinker between actin and microtubule cytoskeletons, offering insights into nuclear movement, a fundamental cellular process.

## Introduction

The nucleus is the largest organelle within a eukaryotic cell, housing what may be its most valuable cargo – the genome. The nucleus is physically coupled to the cytoskeleton via protein networks that extend across the nuclear envelope, establishing direct connections between the surrounding cytoplasm and the nucleoplasm enclosed within it. Linker of nucleoskeleton and cytoskeleton (LINC) complexes are macromolecular assemblies that facilitate nucleocytoskeletal coupling (*1-3*). LINC complexes are formed by single-pass transmembrane Klarsicht/ANC-1/SYNE homology (KASH) proteins (some of which are called nuclear envelope spectrin-repeat proteins (nesprins) in mammals) that span the outer nuclear membrane (ONM). Nesprins contain a C-terminal KASH peptide that directly interacts with inner nuclear membrane (INM) Sad1p/UNC-84 (SUN) proteins within the perinuclear space. SUN proteins interact with nuclear lamins and chromatin within the nucleoplasm and through their interactions with nesprins, establish a direct physical bridge between the nucleoplasmic nuclear lamina and chromatin as well as the cytoplasmic cytoskeleton. This machinery is critical for positioning the nucleus within cells, a function necessary in directionally migrating fibroblasts, developing neurons, and multinucleated muscle cells (*4-6*). It also plays important roles in chromosome pairing during meiosis (*7, 8*). In the nematode *Caenorhabditis elegans*, KASH proteins are required for positioning of nuclei as well as proper organelle positioning within cells (*9-12*). Several human diseases, including ataxias, amyotrophic lateral sclerosis, cardiomyopathies, deafness, and Emery-Dreifuss muscular dystrophy are associated with mutations in nesprin-encoding genes (*13*). These diseases manifest defects in nuclear positioning, highlighting a critical role for nesprins in cellular biology and human health.

Nesprins belong to the spectrin superfamily of cytoskeletal linker proteins that play important biological roles by interacting with the major filamentous cytoskeletal systems within cells, including actin, intermediate filaments, and microtubules. Mammals encode six different nesprins in their genomes, with nesprin-1 and -2 being the most extensively studied. The diverse nesprin family includes members that directly interact with actin filaments (nesprins-1 and -2), and indirectly with intermediate filaments (nesprin-3) and microtubule motor proteins (nesprins-1, -2, and -4) (*14*). Nesprin-1 and -2 are members of a family of giant nesprin proteins that includes *C. elegans* ANC-1 and *Drosophila* MSP-300 (*15*). The largest isoforms of nesprin-1 and -2, encoded by *SYNE1* and *SYNE2*, are 700 KDa and contain tandem N-terminal actin-binding calponin homology (CH) domains, > 50 spectrin repeats (SRs), and a C-terminal W-acidic kinesin-1-binding motif located within an unstructured adaptive domain. However, both nesprin-1 and -2 are spliced into various tissue-specific isoforms (*16*). Nesprin-4 also contains a W-acidic motif that binds directly to the kinesin-1 motor complex (*17*), an interaction which we have previously shown is sufficient to partially relieve the strong autoinhibition of this motor, revealing nesprin-4 to be an activating adapter protein for kinesin-1 (*18*). Similarly, KASH5 acts as an activating adapter protein that relieves the autoinhibition of the cytoplasmic dynein motor during meiosis, demonstrating a strong functional relationship between nesprins and microtubule motor proteins (*15, 16, 19-21*). Consequently, the nesprin family utilizes various mechanisms to interact with the cytoskeleton directly and indirectly, but how these diverse interactions are integrated for nesprin function and nuclear homeostasis remains unclear.

Nesprin-2G (Nesprin-2 giant, N2G hereafter) functions to position nuclei in a variety of cell types. The N2G CH domains bind directly to actin, an interaction important for the assembly of transmembrane actin-associated (TAN) lines, linear arrays of N2G and SUN2 that form along dorsal perinuclear actin cables and drive rearward nuclear positioning in fibroblasts and myoblasts polarizing for directional cell migration (*22-24*). The formin protein formin homology 2 domain-containing 1 (FHOD1) binds to SRs 11 and 12 of N2G, providing a secondary connection to the TAN line-associated dorsal perinuclear actin cables that is important for assembly (*25*). Additionally, SRs 51-53 of N2G interact with the actin-bundling protein fascin and this interaction is important for actin-dependent rearward nuclear positioning in fibroblasts polarizing for directional cell migration (PMID: 27554865). Moreover, N2G’s CH domains are required for its redistribution to the leading front of the nucleus in directionally migrating fibroblasts (*26*). N2G also interacts with the dynein activating adapter bicaudal D cargo adaptor 2 (BicD2), which is important for nuclear movement within migrating neural progenitor cells (*27-29*). As mentioned above, N2G also directly interacts with the kinesin light chain (KLC) via its W-acidic motif, which is composed of the amino acid sequence LEWD. This interaction is important for anchoring the centrosome near the nucleus, and nuclear positioning within multinucleated muscle cells (*30, 31*). W-acidic motifs are found in a multitude of kinesin-1 cargo adapter molecules (*32*), including nesprin-4, where this interaction plays a pivotal role in nuclear positioning in inner ear cells (*17, 33*). Importantly, despite its giant size, mini versions of N2G (mN2G) that contain the N-terminal CH domains, a few SRs, and the C-terminal KASH domain, have been shown to rescue depletion of the endogenous gene in a variety of cell types, revealing the long stretch of SRs within the N2G as not being strictly required for N2G function (*22, 27, 34, 35*). While nesprin-1 and -2 can engage both the actin and microtubule cytoskeletons due to their domain architectures, the possibility of simultaneous cytoskeletal engagement and its physiological relevance remain unclear.

Kinesin-1 (KIF5, ‘kinesin’ hereafter) is a ubiquitous microtubule motor protein that transports numerous cargos towards the plus-ends of microtubules. This heterotetrameric motor is composed of a dimer of kinesin heavy chains (KHCs) containing the motor domains, and two copies of KLCs that bind to cargo molecules via their tetratricopeptide repeat (TPR) domains (*32, 36*). The KLC TPRs bind to short peptide sequences termed W- or Y-acidic motifs often found within unstructured segments of larger proteins termed cargo-adapters, which themselves interact with a variety of different organelles or other cellular cargo (*32, 37, 38*). In the absence of cargo, the kinesin heterotetramer assumes an autoinhibited conformation and does not interact with microtubules (*18, 39-41*). The binding of cargo adapter proteins to the KLC is sufficient to relieve kinesin motor autoinhibition, but robust kinesin motility requires a third component in the binding of a MAP7 family protein to the KHC (*18*). MAP7 proteins are non-enzymatic microtubule-associated proteins (MAPs) that bind both the KHC and microtubules, strongly enhancing the landing rate of kinesin onto microtubules (*18, 42-44*). MAP7 proteins are thought to help kinesin remain bound to microtubules as the motor moves processively along them carrying cargo attached to the KLC within cells (*18, 44*). Hence, the binding of cargo-adapter molecules along with MAP7 synergistically activates the strongly autoinhibited kinesin motor for robust cargo transport along microtubules within cells (*18*). It remains unknown how the binding of cargo-adapter molecules and MAP7 alleviate kinesin autoinhibition.

In this study, we utilized biochemistry and single molecule biophysics to explore the molecular basis of N2G’s interactions with both actin filaments (F-actin) and kinesin. We characterized the dynamic oligomeric states of purified mN2G constructs, providing new insights into how N2G may perform its functions within cells. We performed single molecule measurements of mN2G interactions with F-actin and found that mN2G acts synergistically with MAP7 proteins to relieve kinesin autoinhibition. Our results also show that the binding of MAP7 proteins to kinesin enhances the binding of mN2G to the KLCs, revealing a previously unknown feedback mechanism between MAP7 and kinesin cargo. Finally, we observed that mN2G acts as a cytoskeletal crosslinker by simultaneously interacting with both F-actin and kinesin-1 motors, facilitating kinesin-driven transport of F-actin along microtubules. Our results uncover a novel mechanism for active, motor-driven cytoskeletal crosstalk that may play an important mechanochemical role in nuclear cell biology.

## Results

### Characterization of Recombinant mN2G Constructs and Their Interactions with F-Actin

Human N2G (encoded by the *SYNE2* gene) is a 6885 amino acid long protein with a calculated molecular weight of 782.7 kDa (Fig. 1A), precluding a biochemical analysis of the full-length protein thus far. For this reason, we decided to produce a recombinant construct that encodes mN2G (hereafter: mN2G^SR^), which rescues the nuclear positioning phenotypes observed in tissue culture cells or migrating post-mitotic neurons depleted of endogenous N2G during brain development (*22, 27, 34*). Our mN2G^SR^ construct has a calculated MW of 165.4 kDa and consists of the N-terminal tandem actin-binding CH domains, the first two predicted SRs fused to the C-terminus consisting of five predicted SR repeats with an unstructured adaptive domain containing the KLC-binding W-acidic LEWD motif, and a C-terminal fusion with superfolder green fluorescent protein (sfGFP)-6xHis tag (Figs. 1A and S1A). We removed the C-terminal transmembrane α-helix and KASH peptide (Fig. 1A) to generate a soluble construct for use in solution-based biochemical assays. We used AlphaFold2 (*45*) to predict the structure of mN2G^SR^ (Fig. 1A) The predicted SR54, appears unstructured and contiguous with the unstructured adaptive domain (*46*) containing the LEWD sequence (Fig. S1A). We also generated a smaller construct, mN2G^CH^, that includes only the N-terminal CH domains for use in our assays (Figs. 1A and S1B). Both proteins were purified from bacteria to relative homogeneity (Fig. 1B). We used mass photometry (*47, 48*) to determine the oligomeric state of the purified recombinant mN2G proteins in solution. mN2G^SR^ appeared monomeric (Fig.1C), even across a 20-fold range of concentrations (Fig. S1C). The shorter mN2G^CH^ construct appeared largely dimeric at higher concentrations but transitioned to a monomer at lower concentrations (Figs. 1C and S1C), suggesting that the oligomeric state of N2G is dynamic and dependent on the presence of specific domains within the protein.

**Figure 1:**
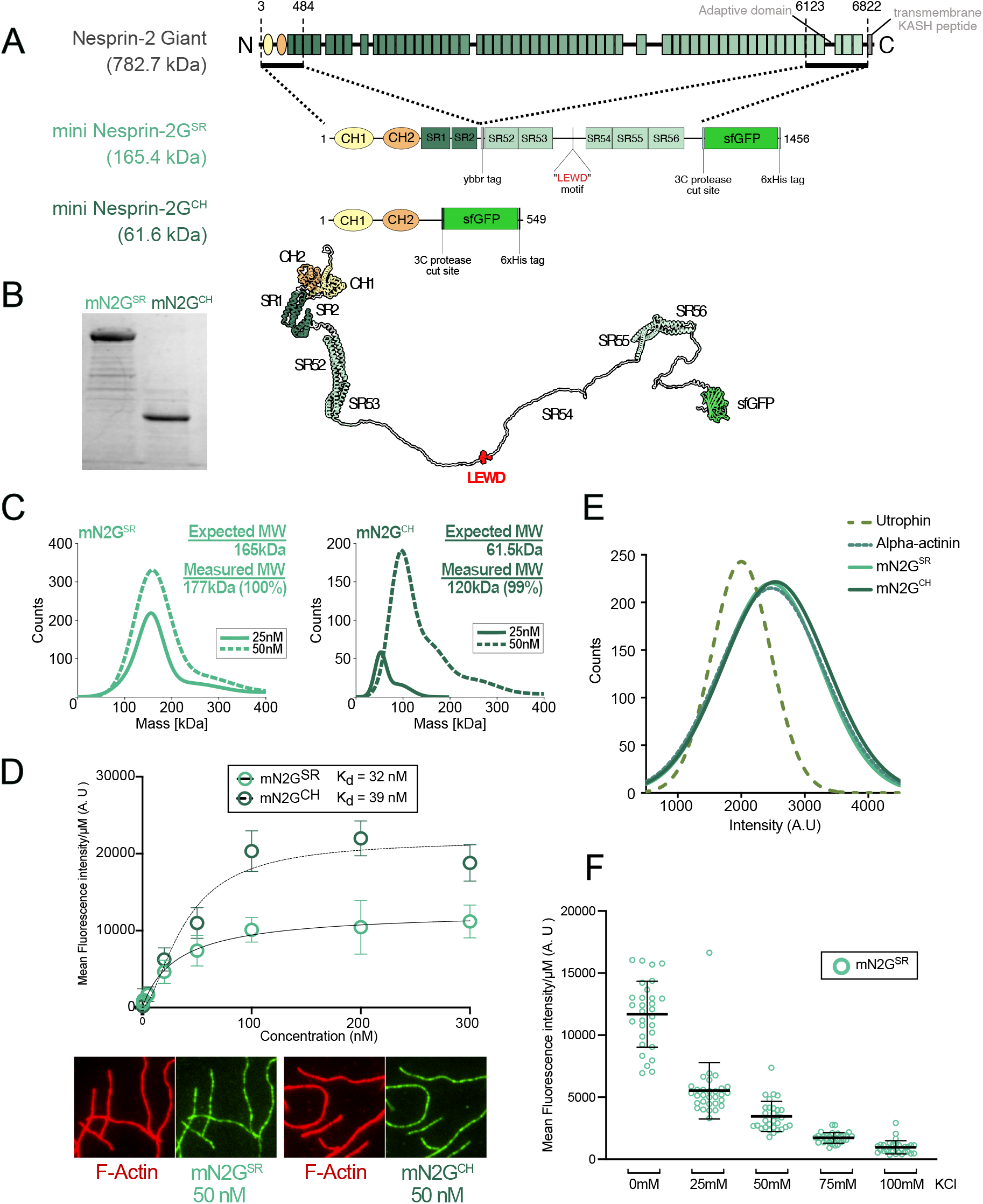
Characterization of recombinant mN2G constructs. (A) A schematic of the native mouse N2G protein as well as the mN2G constructs mN2G^SR^, and mN2G^CH^ used in this study. Schematics highlight protein domains and inserted sequences in the recombinant proteins used in this study. Bottom: AF2 predicted structure of mN2G^SR^. (B) A Coomassie-blue-stained protein gel showing the purified mN2G^SR^ and mN2G^CH^ proteins used in this work. (C) Gaussian fits of the histograms of the measured molecular weight of the indicated proteins in solution obtained using mass photometry at different concentrations. The calculated mass and percentage of particles within the peaks are indicated for each protein. For mN2G^SR^, n = 3931 and 7256 molecules (at 25 and 50nM respectively) and for mN2G^CH^, n = 559 and 3626 molecules (at 25 and 50nM respectively). (D) Top: Mean fluorescence intensity measurements of F-actin bound mN2G^SR^ and mN2G^CH^ (N=2, n=20 actin filaments for each concentration). Bottom: TIRF images of 50nM mN2G^SR^ or mN2G^CH^ binding to phalloidin labeled F-actin. Scale bar: X nm. (E) Gaussian fits of histograms of single-molecule fluorescence intensity distribution for the indicated mN2G proteins compared to other actin-binding proteins bound to F-actin. n = 323, 721, 754, and 271 molecules of utrophin, alpha-actinin, mN2G^SR^, and mN2G^CH^, respectively. (F) Plot of mean fluorescence intensity of mN2G^SR^ bound to F-actin at different concentrations of KCl. N = 2, n = 30 actin filaments measured for each concentration.

To validate the functionality of our recombinant mN2G proteins, we first performed fluorescence-based actin binding assays using total-internal reflection (TIRF) microscopy. Prior bulk biochemical assays have demonstrated that N-terminal fragments of N2G bind directly to actin filaments (*22, 23*). Consistently, both mN2G^SR^ and mN2G^CH^ associated robustly with phalloidin-stabilized and surface-immobilized actin filaments (F-actin, Fig. 1D). Phalloidin stabilization did not affect mN2G’s ability to bind F-actin (Fig. S1D); we thus utilized phalloidin-stabilized F-actin in all subsequent experiments. By quantifying mN2G fluorescence intensity along actin filaments across a range of protein concentrations, we calculated similar apparent affinity constants for both mN2G^SR^ and mN2G^CH^ in the nanomolar range, confirming their high-affinity binding to F-actin (Fig. 1D). We conclude that both mN2G constructs strongly interact with F-actin, confirming the functionality of their CH domains as previously reported (*22, 23*).

Because the oligomeric state of N2G may be dynamic, we sought to determine the oligomeric state of the mN2G constructs bound to F-actin. We utilized two CH domain-containing reference proteins that also bind F-actin with known oligomeric states. Specifically, we purified an N-terminal fragment of utrophin, a monomeric protein (*49*), and full-length α-actinin, a dimer ((*50*) Fig. S1E). We confirmed the expected oligomeric states of these proteins by mass photometry (Fig. S1F). Next, we compared the single molecule brightness of these proteins to our mN2G constructs bound to F-actin. Consistent with the solution mass photometry data (Fig. 1C), the brightness of single mN2G^CH^ molecules was highly similar to the brightness of the dimeric α-actinin protein, while utrophin molecules were significantly dimmer, consistent with their monomeric state (Fig. 1E). Surprisingly, the brightness of mN2G^SR^ molecules closely matched that of α-actinin and mN2G^CH^, indicating that mN2G^SR^ likely dimerizes when bound to F-actin. These results further reveal the dynamic nature of oligomerization of N2G and indicate that F-actin-binding may trigger dimerization of the mN2G^SR^ construct. Finally, the binding of mN2G^SR^ to F-actin is sensitive to ionic strength (Fig. 1F), indicating an electrostatic contribution to their interaction. We conclude that purified mN2G strongly interact with F-actin via ionic interactions, and this interaction facilitates the dimerization of the longer mN2G^SR^ protein.

### Single Molecule Characterization of the mN2G-Actin Interaction

To understand the dynamics of the N2G-actin interaction, we utilized time-lapse single-molecule TIRF imaging. We titrated concentrations of both mN2G constructs, along with utrophin, so we could observe the binding dynamics of single mN2G molecules to individual actin filaments (Fig. 2A). We observed binding and prolonged interactions of single nesprin molecules with F-actin (Fig. 2A). Quantification of interaction times revealed that both mN2G constructs remained bound to F-actin for at least ∼2-fold longer than utrophin (Fig. 2A). We observed a much higher frequency of very short interactions with F-actin for utrophin, as compared to the mN2G constructs (Fig. 2A), contributing to utrophin’s overall shorter F-actin interaction time (Fig. 2A, arrowheads). Quantification of single-molecule landing rates on F-actin revealed that utrophin displayed a ∼20-fold higher landing frequency than mN2G^SR^ (Fig. 2B), consistent with the increased number of short interaction events observed for utrophin (Fig. 2A). The mN2G^SR^ construct showed a reduced interaction time with F-actin as compared to mN2G^CH^ (Fig. 1C, E). Since all these proteins contain tandem CH domains (*51*) with different oligomeric states, the differences in interaction times may be due to dimeric molecules having stronger interactions with F-actin. We also cannot rule out that differences in protein sequence between N2G and utrophin CH domains (52.3% similar, 35.7% identical between human proteins) may also contribute to their differences in F-actin affinity. Nesprin-1 and -2 contain a uniquely longer disordered linker sequence between their two CH domains (*51, 52*), which may play a role in the N2G-F-actin interaction (Figs. 2C, S2). We conclude that mN2G molecules interact with F-actin for many seconds *in vitro*, consistent with their high apparent affinity for F-actin (Fig. 1D).

**Figure 2:**
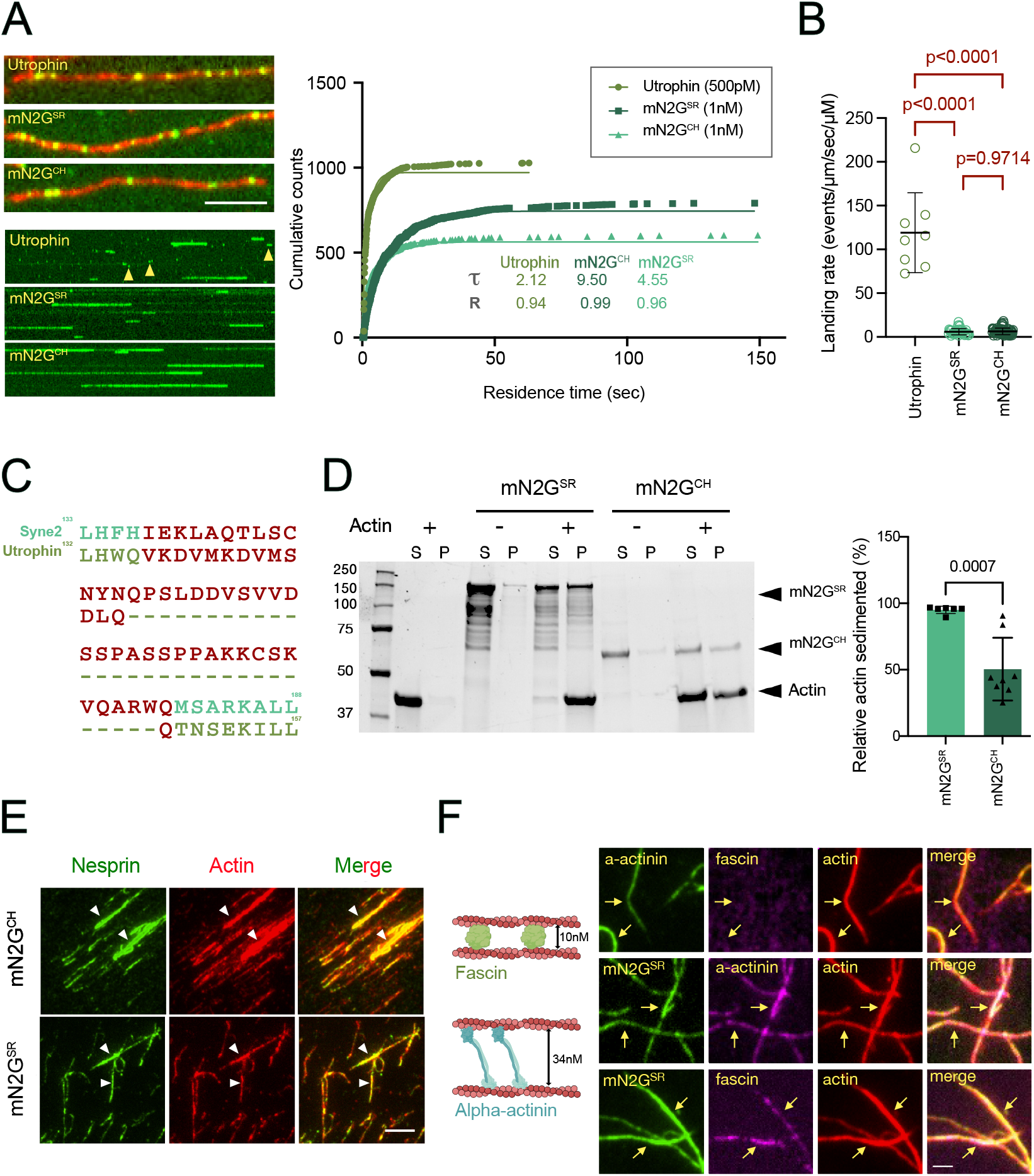
Characterization of recombinant mN2G constructs. (A) TIRF images (left top), kymographs (left bottom), and a cumulative frequency plot (right) depicting the single-molecule binding events of utrophin, mN2G^SR^ and mN2G^CH^ on F-actin. Yellow arrowheads indicate molecules with very short residence times. Calculated residence time constants (τ) and goodness of fit (R) for one phase exponential decay are depicted. A total of 1028 molecules (3 replicates)/606 molecules (4 replicates)/791 molecules (5 replicates) were analyzed for utrophin, mN2G^SR^ and mN2G^CH^ respectively. (B) Quantification of the single-molecule landing rate on F-actin for the indicated constructs. A total of 8 actin filaments (3 replicates)/8 actin filaments (4 replicates)/38 microtubules (5 replicates) were analyzed for utrophin, mN2G^CH^ and mN2G^SR^, respectively. P values: one-way ANOVA. (C) Sequence alignment of the inter-CH domain linker region from N2G and utrophin. See also Fig. S2. (D) Coomassie stained SDS-PAGE gel depicting low-speed co-sedimentation assay. Right: quantification of the band density of actin in the pellet for each condition. P values: two-tailed t-test (n=5). (E) TIRF images depicting that either mN2G^SR^ or mN2G^CH^ (green) can bundle actin filaments (red). Arrowheads indicate actin bundles formed by the indicated mN2G construct. (F) TIRF images of mN2G^SR^ co-assembled into actin bundles with either α-actinin or fascin. Arrows indicate actin bundles decorated by both mN2G and α-actinin or fascin.

In cells, N2G is reorganized into TAN lines along dorsal perinuclear actin cables that are crucial for actin-dependent rearward nuclear positioning during centrosome orientation in fibroblasts and myoblasts polarizing for migration (*22, 53*). Since actin-binding by N2G is required for TAN line formation and subsequently rearward nuclear positioning (*22*), we wondered if N2G alone were sufficient to drive F-actin bundling, a function that could underlie TAN line formation *in vivo*. In solution-based actin bundling assays, we observed that both mN2G^SR^ and mN2G^CH^ induced the pelleting of F-actin bundles by low-speed centrifugation, suggesting the formation of large molecular complexes, as previously reported ((*23*), Fig. 2D). Quantification revealed that mN2G^SR^ drove the bundling of F-actin more efficiently than mN2G^CH^ (Fig. 2D). We examined the protein mixtures by TIRF microscopy and observed large bundles of F-actin and mN2G^SR^, but smaller bundles and more individual F-actin filaments decorated with mN2G^CH^, consistent with the above-described actin bundling results. These results reveal that mN2G^SR^, a construct that rescues N2G function in N2G-depleted fibroblasts polarizing for migration (*22*) and in migrating post-mitotic neurons during brain development (*27*), can induce the formation of large F-actin bundles *in vitro*. The reason for the higher bundling efficiency of mN2G^SR^ is unclear but may be related to the more dynamic oligomeric state of mN2G^CH^ relative to mN2G^SR^ (Fig. 1B, S1C) or its longer F-actin interaction times (Fig. 2A), which may hamper its bundling efficiency.

F-Actin bundling is a common function of actin binding proteins that harbor distinct types of molecular architectures. For instance, N2G’s tandem CH domains followed by SRs is an architecture similar to what is observed for the actin-crosslinking protein a-actinin (*51*). However, a-actinin forms an extended anti-parallel dimer that mediates F-actin crosslinking with a spacing of 34 nM (*54*), whereas the structure of mN2G^SR^ dimer bound to F-actinin our assays is unknown (Fig. 1E). In contrast, fascin is a monomeric actin crosslinking protein with a distinct compact shape that mediates tight F-actin bundling with a spacing of ∼10 nm (*55, 56*). Prior work found that fascin and a-actinin exclude each other within F-actin bundles due to their distinct protein architectures (Fig. 2F) (*54*). Having established N2G as another F-actin-bundling/crosslinking factor (Fig. 2E-F), we wondered if the N2G-F-actin interaction were affected by the presence of distinct actin-crosslinkers in the same system. Purified a-actinin and fascin behaved as dimers or monomers respectively in solution, as expected (Fig. S1F). When combined with F-actin, fascin was largely excluded from a-actinin positive F-actin bundles, similar to prior results (Fig. 2F) (*54*). When a-actinin or fascin were mixed with mN2G^SR^ and F-actin, we observed robust mN2G^SR^ binding to F-actin bundles positive for either protein, suggesting that mN2G^SR^ is not excluded from F-actin bundles in the presence of distinct F-actin-crosslinkers (Fig. 2F). We conclude that mN2G^SR^ is a robust F-actin crosslinker that is not sensitive to the presence of distinct F-actin crosslinkers or actin bundle architectures. We speculate that this feature could be important for N2G’s role in TAN line formation *in vivo*.

### N2G Connects Kinesin-1 To F-actin via the KLC in a MAP7D3-Dependent Manner

Having established N2G’s robust interaction with F-actin, we focused on N2G’s reported interaction with the microtubule motor kinesin-1 (*31*). N2G’s C-terminal adaptive domain contains the W-acidic motif that binds the KLC TPR domain (*31, 37, 38*), but this interaction has not been investigated in a biochemically pure system. Given the physical separation of the N-terminal actin-binding CH domains and C-terminal W-acidic KLC-binding motif (Fig. 1A), we reasoned it may be possible for N2G to bind to both F-actin and kinesin simultaneously. Recombinant kinesin heterotetramers composed of the KIF5B KHC and KLC2 (K5B2) did not directly interact with immobilized F-actin (Fig. 3A), nor with F-actin bound to mN2G^CH^, which lacks the W-acidic motif. In contrast, in the presence of mN2G^SR^, we observed clear recruitment of kinesin tetramer to F-actin (Fig. 3A, B). With mN2G^SR^, kinesin recruitment to F-actin was enhanced over ∼115-fold as compared to mN2G^CH^ (Fig. 3B), revealing that mN2G^SR^ is capable of simultaneously interacting with both F-actin and kinesin.

**Figure 3:**
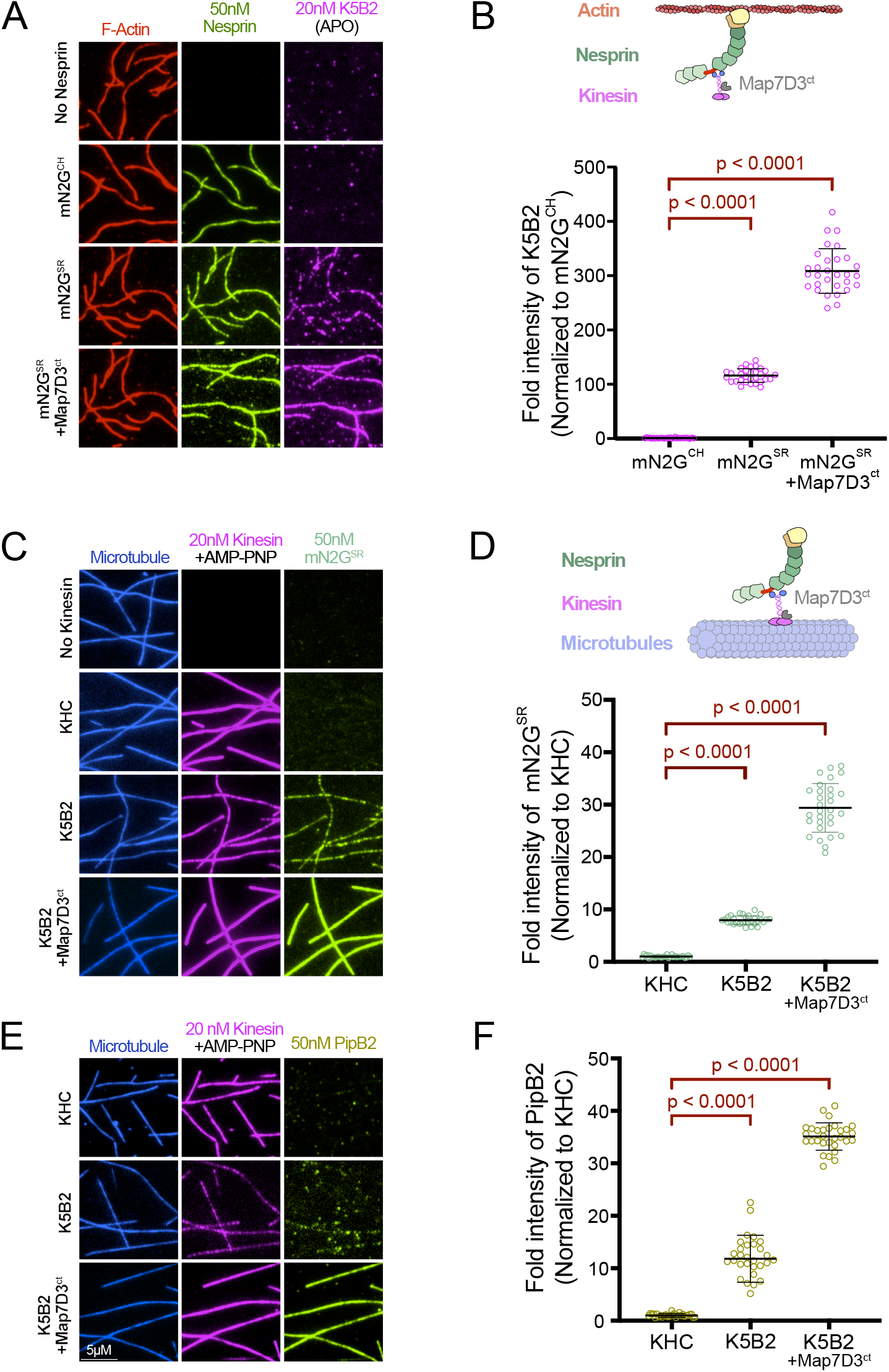
MAP7D3 Facilitates the Kinesin-mN2G Interaction in the Absence of Microtubules. (A) Representative TIRF images showing the recruitment of the indicated mN2G constructs and kinesin heterotetramers (KIF5B/KLC2:K5B2, magenta) to actin filaments (red) and the effects of MAP7D3^ct^ on this interaction. (B) Top: A schematic showing the mN2G-kinesin-MAP7D3^ct^ complex interacting with actin filaments. Bottom: quantification of the fold intensity (normalized to the mN2G signal present on actin filaments) for each condition. P Values: one-way ANOVA followed by Dunnet’s test. N=2, n=30 actin filaments quantified for each concentration. (C) Representative TIRF images of the recruitment showing recruitment of mN2G^SR^ (green) onto microtubules (blue) coated in indicated kinesin constructs (magenta) in the rigor state (AMP-PNP), and the effects of MAP7D3^ct^ on this interaction. (D) Top: schematic showing the mN2G^SR^-kinesin-MAP7D3^ct^ complex interacting with microtubules. Quantification of the fold intensity (normalized to the amount of kinesin signal) for each condition. P values: One-way ANOVA followed by Dunnet’s test. N=2, n=30 actin filaments for each concentration. (E) Representative TIRF images of the recruitment of PipB2 (olive) to microtubules (blue) in the presence of the indicated kinesin constructs, and the effects of MAP7D3^ct^ on this interaction. (F) Quantification of the fold intensity (normalized to the amount of kinesin signal) for each condition. P values: one-way ANOVA followed by Dunnet’s test. N=2, n=30 actin filaments for each concentration.

The kinesin heterotetramer is strongly autoinhibited through intramolecular folding that prevents its ability to interact with microtubules (*18, 39, 40*). The binding of cargo adapter molecules to the KLC, and MAP7 proteins to the KHC is thought to be required to fully relieve kinesin autoinhibition (*18, 40*). It is unknown if these interactions feedback to one another within the context of the intact kinesin heterotetramer, or if microtubule-binding by the motor alters these interactions. We therefore asked if the binding of MAP7 proteins to the KHC would affect the interaction of the KLC with mN2G^SR^ on F-actin in the absence of microtubules. We included in our assay the C-terminal kinesin-binding domain of MAP7D3 (MAP7D3^ct^), a protein that interacts directly with the KHC *in vitro*, and is important for kinesin function *in vivo* (*43*). Strikingly, the inclusion of MAP7D3^ct^ resulted in a ∼3-fold enhancement of kinesin recruitment to F-actin coated with mN2G^SR^ (Fig. 3A, B). This result was not anticipated a priori, as MAP7 interacts with the KHC (*42, 57*), while cargo adapter molecules such as N2G bind to the KLC (*31, 37*). In the absence of mN2G^SR^, MAP7D3^ct^ alone did not facilitate kinesin binding to F-actin (Fig. S3). Hence, the binding of kinesin cargo to the KLC is strongly enhanced by MAP7 proteins that interact with the KHC, even in the absence of microtubules. These results also demonstrate that N2G facilitates the recruitment of kinesin homotetramer onto F-actin, establishing a direct connection between the two major cytoskeletal systems. They also reveal a previously unknown feedback mechanism between MAP7 binding to the KHC and cargo molecules binding to the KLC, implying intramolecular regulation between the cargo- and microtubule-binding portions of the kinesin heterotetramer.

Next, we sought to define the interaction between N2G and kinesin in the context of microtubule binding. Purified mN2G^SR^ did not interact with surface-immobilized, taxol-stabilized microtubules in the absence of kinesin, nor in the presence of immobilized KHC bound to microtubules (Figs. 3C-D). In contrast, microtubule-bound kinesin heterotetramer clearly recruited mN2G^SR^ to microtubules (Figs. 3C-D). Inclusion of MAP7D3^ct^ again resulted in strong further enhancement of kinesin binding to microtubules (Fig. 3C), as expected from prior work (*42, 43*). MAP7D3^ct^ also facilitated a large increase in the amount of mN2G^SR^ that bound to kinesin (Figs. 3C-D), reinforcing our observation that MAP7D3^ct^ also stimulates the binding of F-actin-bound mN2G^SR^ to the KLC (Fig. 3A, D). These findings further show that N2G can recruit kinesin to F-actin and reveal a previously unknown intramolecular feedback mechanism whereby MAP7 protein binding to the KHC augments the binding of cargo-adapter molecules to the KLC independently of microtubules.

Finally, we wondered if the effect of MAP7D3^ct^ on kinesin cargo-adapter binding was specific to mN2G, or if it applied more broadly to various types of kinesin cargo-adapters. We repeated the microtubule recruitment assay with an orthogonal type of kinesin cargo-adapter: the *Salmonella* effector protein PipB2, which recruits host kinesin molecules to the intracellular *Salmonella* vacuole (*58*). PipB2 contains a C-terminal Y-acidic motif which binds to the KLC TPR domains in a distinct manner compared to the W-acidic motif found in N2G (*32, 37, 38*). We observed that microtubule-bound kinesin dimers lacking the KLC did not recruit PipB2 to microtubules (Figs. 3E-F and S1E). Microtubule-bound kinesin tetramers bound relatively poorly to PipB2, possibly due to KLC autoinhibition, or cargo-adapter autoinhibition, a common theme emerging in molecular motor activation (*18, 39, 40, 59*). Once again, the inclusion of MAP7D3^ct^ strongly enhanced the binding of PipB2 to microtubule-bound kinesin heterotetramers (Fig. 3E, F). These results further confirm that MAP7D3^ct^ facilitates the binding of multiple types kinesin cargo-adapters containing either W- or Y-acidic motifs, and reveals that this principle applies to kinesin cargo-proteins that span the kingdoms of life. The relative differences in the magnitudes of cargo-adapter binding suggest that further regulatory mechanisms may govern the specific affinities of distinct cargo-adapters to the KLC and this point requires further efforts to dissect.

### Active Cytoskeletal Crosslinking via the N2G-Kinesin Complex

Having established that mN2G^SR^ can simultaneously interact with both F-actin and kinesin, we wondered if it were possible that this complex could directly link F-actin to microtubules via the kinesin motor (Fig. 4A). Further, this linkage is expected to be dynamic in the presence of ATP, as the kinesin motor cycles through its strong and weak microtubule-binding states (*60*). We mixed kinesin, mN2G^SR^, and F-actin together and introduced them into a chamber containing surface-immobilized microtubules. In the presence of ATP, we did not observe obvious recruitment of F-actin to microtubules, possibly due to the relatively low landing rate of cargo-adapter bound kinesin on microtubules in the absence of MAP7 proteins. Notably, when MAP7D3^ct^ was included, we observed recruitment and alignment of F-actin along microtubules in the presence of ATP (Fig. 4B, Movie S1). Using multi-channel TIRF imaging, we confirmed the presence of both kinesin and mN2G^SR^ enriched with the F-actin along microtubules (Fig. 4B, Movie S1), confirming that the kinesin-mN2G^SR^ co-complex was responsible for F-actin recruitment to microtubules. Thus, N2G crosslinks F-actin to microtubules via the kinesin motor in a MAP7D3-dependent manner.

**Figure 4:**
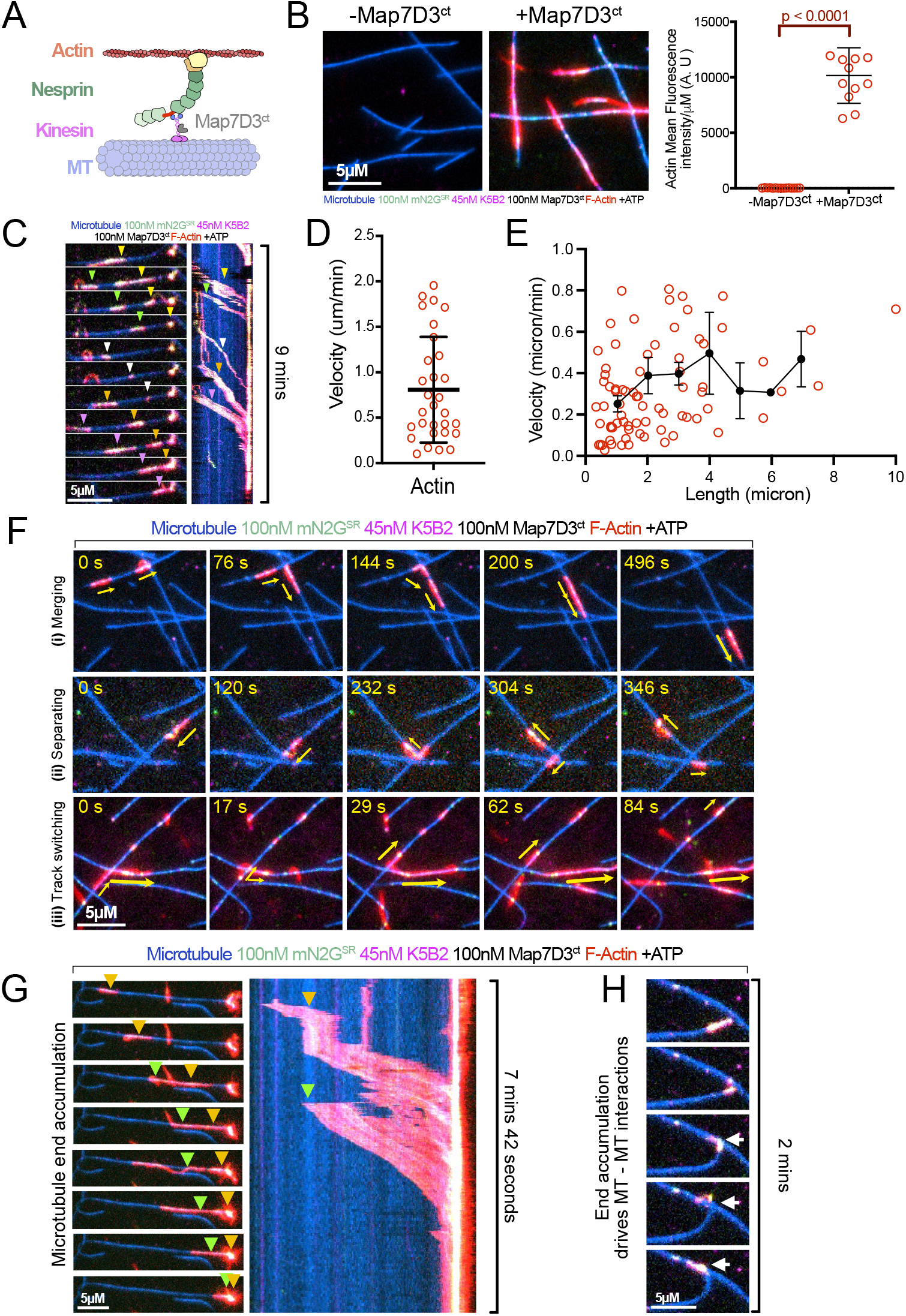
Active Actin-microtubule crosstalk via the mN2G^SR^-kinesin-MAP7D3^ct^ complex. (A) schematic showing the mN2G^SR^-kinesin-MAP7D3^ct^ complex interacting with both microtubules and actin filaments. (B) Representative TIRF images of actin recruitment on surface immobilized microtubules via mN2G^SR^-K5B2 with or without MAP7D3^ct^. Quantification of mean fluorescence density of actin on microtubules with or without MAP7D3^ct^. N=2, n=30 actin filaments for without MAP7D3^ct^ and N=3, n=8 to 12 actin filaments for with MAP7D3^ct^. P values: student’s two-tailed t-test. (C) Still frames of a representative TIRF movie of actin along with the mN2G^SR^-kinesin-MAP7D3^ct^ complex moving along microtubules in the presence of ATP. Arrowheads denote individual actin filaments as they land and are transported along the microtubule. Right: kymograph of the entire time series showing multiple F-actin filaments transported along the microtubule. Note the accumulation of F-actin at the microtubule end. (D) Quantification of the velocity of actin filaments moving on microtubules. N=3, cumulative n=30 actin filaments across all experiments. (E) A plot of the velocity of actin filaments vs. their lengths. N=12, cumulative n=75 across all experiments. We have grouped the velocity data points into intervals of 1 unit length (0-1, 1-2…) and then plotted the trend with mean and error (black). (F) Examples of (i) merging, (ii) separating, and (iii) track switching events of actin behaviors during their movement along microtubules. (G) Representative images of the transport, merging and accumulation of actin-mN2G^SR^-kinesin-MAP7D3^ct^ complexes at microtubule ends. Arrowheads denote individual actin filaments as they land and are transported along the microtubule. (H) Representative images of an actin-mN2G^SR^-kinesin-MAP7D3^ct^ complex at a microtubule end bridging between two microtubules and causing the bending and reorientation of one microtubule relative to the other.

Next, we utilized multi-channel time-lapse TIRF imaging to reveal the dynamics of the interactions between F-actin and microtubules driven by the mN2G^SR^-kinesin -MAP7D3^ct^ co-complex. Remarkably, we observed persistent unidirectional transport of single actin filaments and bundles along microtubules (Fig. 4C), revealing that mN2G^SR^ facilitates the dynamic bundling and crosslinking of actin and microtubules driven by kinesin motors *in vitro*. Actin filaments were actively co-aligned with the microtubules on which they translocated. Fluorescent signals for kinesin and mN2G^SR^ were highly enriched on microtubule-associated actin filaments (Fig. 4C), suggesting that a high density of kinesin-mN2G^SR^ complexes underlies the movement of F-actin along microtubules. Consistently, the velocity of F-actin transport along microtubules was much slower than single kinesin movement (*18*), at ∼ 0.8 μm/min (Fig.4D). We speculate the relatively slow velocity of F-actin movement may be due to the high density of kinesin motors bound to F-actin resulting in interference of kinesin stepping dynamics. We measured the velocity of actin filaments as a function of their length and found that there was not a strong correlation of F-actin velocity with the length of a motile actin filament (Fig. 4E). These results reveal that a N2G-kinesin-MAP7D3 system drives the crosslinking and active remodeling of an F-actin-microtubule cytoskeletal composite *in vitro*.

We noticed that F-actin movement along microtubules was highly processive. Due to this persistence, transported actin filaments displayed diverse behaviors as they moved along the microtubule surface. We observed actin filaments merging into larger bundles during their transport along microtubules (Fig. 4F (i), Movie, S2). Actin filaments are sensitive to mechanical breakage (*61*) and we frequently observed actin filaments and bundles fragmenting when different regions of an actin filament were bound to distinct microtubules and pulled in opposite directions (Fig. 4F (ii), Movie S2). When a single actin filament reached a microtubule-microtubule intersection, we often observed track switching of the leading portion of the actin filament onto the crossing microtubule (Fig. 4F (iii), Movie S2). Because of the high processivity of the transported actin filaments, many merged with other filament bundles along the same microtubule and moved to the microtubule end where they were often retained for the remainder of the imaging period (Fig. 4C, G, Movie S3). We noticed that these clusters of actin bundles at the ends of microtubules could interact with nearby microtubules, creating a dynamic bridge between them. This interaction caused one microtubule to bend and align with the other (Fig. 4G, Movie S3). We conclude that mN2G is capable of underpinning dynamic cytoskeletal crosstalk between microtubules and actin via its strong binding to F-actin and activation of autoinhibited kinesin in conjunction with MAP7D3.

## Discussion

In this study, we have reconstituted a unique molecular system that directly connects two of the most prominent cytoskeletal filaments found in cells, actin and microtubules, facilitating the ATP-dependent active remodeling of the filament composite driven by the kinesin-1 motor. Central to this system is N2G, a giant KASH protein that plays a myriad of roles in the cell biology of the nucleus. N2G acts as both a static anchor to F-actin via its N-terminal CH domains, and an activating cargo-adapter for kinesin via its C-terminal W-acidic motif. We found that the tandem CH domains of N2G have uniquely prolonged interactions with F-actin and crosslink F-actin into bundles. Further, we found that N2G is capable of binding F-actin bundles formed by actin crosslinking proteins with distinct molecular architectures, suggesting that the N2G-F-actin interaction is robust against the molecular composition of actin bundles in cells. These findings shed light on the potential mechanisms underlying the formation of TAN lines in cells, whereby N2G is reorganized into linear arrays in response to the formation of dorsal perinuclear actin cables in fibroblasts polarizing for migration through its prolonged F-actin interactions and insensitivity to local actin architecture. In conjunction with its direct interaction with the F-actin bundling formin FHOD1 and fascin (*25, 62, 63*), N2G appears well suited to drive the formation of higher-order actin structures such as TAN lines.

Within the native N2G protein, the CH domains and kinesin-binding W-acidic motif are separated by ∼6000 amino acids, suggesting that a considerable nanoscale distance may exist between the F-actin and microtubules crosslinked by N2G-kinesin complexes. Nevertheless, our results generated with mN2G^SR^ demonstrate that a large distance between the actin and microtubule binding domains is not required for functional crosslinking via the kinesin motor, itself a highly elongated molecule of ∼ 70-80 nm (*39, 40*). Additionally, we found that MAP7 proteins play a critical role in this process by strongly augmenting kinesin binding to N2G’s W-acidic motif and microtubules, consistent with growing evidence that MAP7 protein function is critical for kinesin-1 activity within cells (*43, 64, 65*). Our reconstitution reveals a novel role for N2G in facilitating an active motor-driven connection between the actin and microtubule cytoskeletons and reveals new molecular features that further enlighten the mechanisms of nesprins in cellular biology.

We found that the F-actin-binding dynamics of N2G differ significantly from those observed for other tandem CH domain-containing actin-binding proteins. Our data suggest that N2G molecules in solution adopt a configuration that biases the molecule towards a monomeric state, and that F-actin-binding alters this configuration, stimulating dimerization when bound to F-actin. Truncation of the SRs adjacent to N2G’s CH domains facilitated the concentration-dependent dimerization of mN2G^CH^ in solution, suggesting that these SRs may regulate the oligomeric state of giant KASH proteins. In contrast, the utrophin N-terminus is monomeric in solution and when bound to actin, likely contributing to its lower residence time on F-actin. These differences may further stem from sequence differences present within the tandem CH domains, and possibly the extended linker segment found between the giant KASH protein CH domains (*52*). Further work is necessary to understand the regulatory mechanisms that control N2G oligomerization and function, but our results suggest that F-actin binding relieves an autoregulatory conformation that controls N2G oligomerization.

The prevailing model of LINC complexes is that a trimer of SUN proteins interacts with three KASH proteins in the perinuclear space. Structures predict that the three KASH proteins cross the outer nuclear membrane independently at distances too far for their trans-membrane spans to interact (*66*). The oligomeric state of the cytoplasmic domains of KASH proteins are poorly understood. This absence of trimerization could be due to the lack of SUN proteins in our experiments, or, reflect a model where cytoplasmic domains of KASH proteins are free to oligomerize independently of the three C-terminal KASH peptides interacting with a SUN protein trimer within perinuclear space.

The cytosolic domain of the meiotic KASH protein KASH5 exists predominantly as a dimer due to the presence of a central coiled-coil (*20, 21*). Dimeric KASH5 activates the dimeric autoinhibited dynein motor complex, similar to our observations with nesprin-4 (*18*) and N2G with the dimeric autoinhibited kinesin motor in this study. Hence, it seems plausible that dimeric KASH proteins align with the structural symmetry of dimeric microtubule motor complexes with which they engage. We postulate that a trimeric nesprin-SUN complex that interacts with a dimeric kinesin motor would result in one unoccupied N2G subunit. This could conceivably promote the formation of higher-order N2G-SUN1/2-kinesin oligomers through polymerization of adjacent kinesin-nesprin complexes. Recent structural data obtained using truncated SUN protein luminal domains bound to KASH peptides from various nesprins indicate higher-order oligomerization via SUN trimer interactions (*3, 67, 68*). Therefore, more complex oligomerzation of both the luminal KASH-SUN complex, as well as potentially the KASH protein cytosolic domains bound to molecular motors, or other effectors, may be a common feature of LINC complex function. Investigating the structural organization of N2G-SUN1/2-motor complexes is crucial to address these intriguing aspects further.

While N2G plays a central role in the actin-microtubule crosstalk we observe, our data also illuminate a previously unknown role for MAP7 proteins in facilitating the KLC-nesprin interaction. MAP7 proteins have emerged as critical kinesin regulatory molecules that bind directly to the KHC (*18, 42, 43*), dramatically enhancing its landing rate onto microtubules, and potentially facilitating its release from the autoinhibited conformation termed the “lambda” conformation (*39, 40*). However, a direct role for MAP7 proteins in facilitating kinesin-cargo interactions has not been reported to our knowledge. Our data reveals this role for MAP7D3^ct^, which dramatically enhances the interaction between the KLC and mN2G^SR^, even in the absence of microtubule binding by the KHC. MAP7D3^ct^ is essential for the robust interaction between kinesin and mN2G^SR^ that underlies the active microtubule-actin crosstalk reported here. We utilized a C-terminal fragment of MAP7D3 (MAP7D3^ct^), which binds strongly to the KHC, and rescues kinesin-microtubule binding in cells depleted of all MAP7 proteins (*43*). However, cells express multiple isoforms of MAP7 proteins, which may have distinct roles in regulating kinesin activity, and more work is needed to delineate the specific cellular roles performed by each MAP7 isoform.

Our findings show that MAP7 proteins not only enhance the KHC-microtubule interaction as previously reported (*42, 43*), but also facilitate the interaction of the KLC with distinct cargo molecules. We hypothesize that this arises from MAP7-induced dissociation of the KLC TPR domains from KHC coiled-coils in the autoinhibited lamda conformation of the kinesin heterotetramer (*39, 40*). We found that MAP7 enhances the binding of both W- and Y-acidic KLC-binding motifs present in kinesin cargo, suggesting a fundamental principal for kinesin-cargo interactions. Indeed, even pathogenic proteins utilized to hijack kinesin motility by bacterial pathogens rely on MAP7-enhanced binding to the KLC (*58*), demonstrating the conservation of the role of MAP7 proteins with respect to kinesin cargos that span kingdoms of life. Consequently, we propose that MAP7 proteins govern kinesin-1-based motility within cells through their dual activities of enhancement of KHC-microtubule and KLC-cargo interactions, likely through the simultaneous direct interactions with microtubules and disruption of the autoinhibited lamda kinesin conformation.

What role might active crosslinking between actin and microtubules play *in vivo*? Prior genetic and cell biological evidence suggests complementary, although not always mutually necessary, roles for actin- and microtubule-binding in giant KASH protein biology. First, nuclear movement via TAN lines does not appear to require microtubules in fibroblasts (*22*), whereas nesprin interactions with the microtubule motors kinesin-1 and dynein are critical for nuclear movement within myotubes and during *C. elegans* development (*31, 57, 69, 70*). In *Drosophila* skeletal muscle, genetic experiments revealed direct roles for kinesin, MAP7, and nesprins in nuclear positioning, highlighting a role for MAP7 in nesprin biology that can be explained by our current *in vitro* findings (*57*). Therefore, several lines of evidence reveal roles for both actin and microtubules in nesprin biology, but a direct connection between these two filament systems remains complicated to dissect *in vivo* and has not been rigorously tested to date. The active crosslinking of the two cytoskeletal systems we reveal here suggests a more complicated dynamic interaction between the two major cytoskeletal systems may be at play *in vivo*. These results potentially provide new insights into how actomyosin tension is communicated across the nuclear envelope as revealed by FRET-based nesprin tension biosensors (*71*). Our findings present new assays with which to address the structural organization and function of giant KASH protein-cytoskeletal complexes and suggests further exploration of the cell biological roles of active actin-microtubule crosstalk mediated by giant KASH protein-motor complexes *in vivo*.

## Acknowledgements

We thank all the members of the MOM and Starr/Luxton labs for their continual input and feedback on this project. This work was supported by grants from NIGMS 1R35GM124889 to R.J.M. and 1R35GM133688 to K.M.O.M. Research in the Starr-Luxton lab is supported by NIGMS grant R35GM134859 and an Allen Distinguished Investigator Award, a Paul G. Allen Frontiers Group advised grant of the Paul G. Allen Family Foundation. R.J.M. and K.M.O.M. thank members of the FOM for their continued support and guidance.

## Author contributions

N.S.: conceptualization, investigation, verification, formal analysis, visualization, writing and editing. K.O.: conceptualization, investigation, formal analysis, visualization, editing. A.K. investigation. D.E. formal analysis, visualization, editing. D.S. funding acquisition, conceptualization, editing. K.M.O: funding acquisition, conceptualization, editing. G.W.L. funding acquisition, conceptualization, writing and editing. R.J.M.: funding acquisition, conceptualization, visualization, writing and editing, project administration, verification, methodology.

## Competing interest statement

The authors declare no competing interests.

## Materials and Methods

### DNA constructs

To generate mN2G^SR^, a gBlock (Integrated DNA Technology (IDT), Coralville, IA) encoding amino acids 3-484 fused to aa 612-6822 of mouse N2G that was codon-optimized for expression in E. coli was cloned it into a pet28a vector containing a sfGFP-6xHIS tag. The mN2G^CH^ construct was generated from the previously described mN2G^SR^ by PCR amplifying the relevant cDNA from the previously made mN2G^SR^. To generate Map7D3^ct^, a cDNA fragment encoding aa 426-876 of human Map7D3 was codon-optimized for E. coli and synthesized by gBlocks (Integrated DNA Technologies, Coralville, IA, USA) and incorporated into a pet28a vector containing and a N-terminal StrepII tag. A plasmid encoding human KIF5B-KLC2 plasmid was used as previously described (*18*). A plasmid encoding aa 1-261 of human utrophin (PaGFP-UtrCH, Addgene plasmid #26738) was obtained as a gift from Dr. William Bement (University of Wisconsin, Madison). The cDNA sequence encoding aa 1-261 of human utrophin was PCR amplified and cloned into the pet28a-sfGFP-6xHis by Gibson assembly. We obtained pGEX4T-3-human Fascin (Fscn1, Uniprot ID: Q16658) and pET30a-human α-actinin (ACTN4, Uniprot ID: O43707) as gifts from Dr. David D. Kovar (University of Chicago). The Thrombin cleavage site present in pGEX4T-3 was replaced with a Prescission protease site by Gibson assembly. The entire coding sequence of ACTN4 was PCR amplified and cloned into pET28a-sfGFP-6xhis by Gibson assembly. All constructs were verified by Sanger or Nanopore sequencing.

### Protein expression and purification

Bacterial expression and preparation of mN2G^SR^, mN2G^CH^, Map7D3^ct^, Utrophin, Fascin, and α-actinin were performed as described below. Briefly, BL21-CodonPlus (DE3)-RIPL E. Coli (Agilent, Santa Clara, CA) were transformed with each plasmid and grown at 37ºC in Luria Broth with 50 μg/ml Kanamycin until an optical density at 600 nm of 0.6. The cultures were allowed to return to room temperature and then induced by the addition of 0.2 mM isopropyl-b-D-thiogalactoside overnight at 18ºC with shaking. Bacteria were subsequently harvested by centrifugation at 22,769 xg for 15 min, flash frozen in liquid nitrogen, and stored at -80ºC.

For recombinant protein purification, frozen pellets of bacteria were thawed on ice and resuspended in purification buffer (PB: 50 mM Tris-HCl pH 8.0, 150 mM KCH3COO, 2 mM MgSO_4_, 1 mM EGTA, and 5% glycerol) freshly supplemented with 1 mM phenylmethylsulfonyl fluoride (PMSF), 0.1 mM ATP, NucA nuclease, and protease inhibitor mix (Promega, Madison, WI). For purifying Map7D3^ct^ and fascin 1 mM DTT was also added during this step. For purifying utrophin, fascin, and α-actinin, a buffer containing 50 mM Na-PO4 pH8.0, 150 mM NaCl, 2 mM EGTA, 20 mM imidazole was used instead of PB. Bacteria were lysed by passage through an Emulsiflex C3 high-pressure homogenizer (Avestin, Ottawa, ON, Canada) and the subsequent by addition of 1% Triton X-100 for 5 min on ice. Lysed cells were then centrifuged at 22,769 xg for 20 min at 4° C. For affinity purification, the clarified lysates were incubated with resin as follows. Map7D3^ct^ was incubated with Streptactin XT resin (IBA Lifesciences, Göttingen, Germany) for 1 hr at 4° C and washed with PB buffer. Protein was subsequently eluted with elution buffer (EB: PB buffer plus 50 mM biotin, pH 8.0). mN2G proteins were loaded onto a 5 ml HisTrap HP (Cytiva, Wilmington, DE) at 4° C, washed with PB containing 20 mM immidazole, and eluted with a gradient of 20-300 mM Imidazole in PB. Fractions containing recombinant protein were isolated and frozen at -80° C. Lysates containing fascin were loaded onto a column packed with glutathione Sepharose 4 FF resin (GE Healthcare, Chicago, IL). Fascin recovered from the glutathione column was cleaved with Prescission protease overnight at 4º C. Lysates containing Utrophin or α-actinin were loaded on a column packed with His-Pur Ni-NTA resin (ThermoFisher Scientific, Waltham, MA). The column was washed with 0.6 M NaCl, and bound proteins were eluted with 300 mM Imidazole.All proteins were further purified by gel filtration or ion exchange chromatography.

While mN2G^SR^ fractions were injected directly onto a HiLoad 16/600 Superose 6 pg column (Cytiva), mN2G^CH^ was concentrated down to 500 μl using an Amicon Ultra 5 centrifugal filters (Merck, Darmdstad, Germany) and then injected onto a Superdex 200 10/300 GL column (Sigma-Aldritch, Burlington, MA) in GF150 buffer (25 mM HEPES-KOH pH 7.4, 150 mM KCl, and 1 mM MgCl_2_). Map7D3^ct^ was further purified using a HiTrap Capto S cation exchange chromatography column (Cytiva) equilibrated in HB buffer (35 mM PIPES-KOH pH 7.2, 1 mM MgSO_4_, 0.2 mM EGTA, and 0.1 mM EDTA, pH 7.1). Bound proteins were eluted with a 45 mL linear gradient of 0-1 M KCl in the HB buffer. Cleaved Fascin was concentrated in an Amicon-Ultra MWCO 50 kDa concentrator (Millipore Sigma, Burlington, MA), and loaded onto Phenomenex BioSep SEC-s4000 300 x 21.20 mm column (Phenomenex, Torrance, CA) equilibrated with GF150. Utrophin or α-actinin eluted from a Ni-NTA column (ThermoFisher Scientific) was diluted 3 times and loaded onto 5 ml HiTrap Q HP column (Cytiva). The protein was then eluted with a linear gradient of NaCl from 0 to 600 mM. Peak fractions were pooled and loaded onto a size exclusion column equilibrated with GF150. A Phenomenex BioSep SEC-s4000 300 x 21.20 mm column (Phenomenex) was used for α-actinin, while a Superose 6 Increase column (Cytiva) was used for Utrophin-sfGFP-6xhis and α-actinin-sfGFP-6xHis.

Fascin and α-actinin were labeled with 3M excess 6-TAMRA Maleimide (Aat Bioquest, Pleasanton, CA). The dye was diluted to 10 mM in DMSO, added to the proteins at an excess of 3 molar, and incubated overnight at 4 ºC. Free dye was removed by passing through a 5ml HiTrap desalting column (Cytiva).

Protein-containing fractions were pooled and supplemented with 5% Glycerol. Protein aliquots were subsequently flash frozen by liquid nitrogen and stored at -80°C. Protein concentrations were measured using a NanoDrop One (ThermoFisher Scientific) based on the absorbance of the relevant fluorophores.

KIF55B-KLC2 was purified as previously described (*18*).

### Single-molecule mass photometry assay

Single-molecule mass photometry was carried out as previously described (*72*). Briefly, imaging chambers were prepared by cleaning #1.5 24 x 50 mm Deckgläser glass coverslips (ThermoFisher Scientific) by sonication for 1 hr followed by their storage in isopropanol. Imaging chambers were assembled the day mass photometry was performed. The coverslips were rinsed with fresh isopropanol followed by Milli-Q water and dried with filtered air. CultureWell silicone gaskets (Grace Bio-Labs, Bend, OR) were placed onto the freshly cleaned coverslips providing six independent sample chambers. Samples and standard proteins (Apoferritin, Bovine serum albumin (BSA), and Thyoglobulin, Sigma-Aldrich) were diluted to 10-50 nM in freshly 0.1 μm filtered SRP90 buffer (90 mM HEPES-KOH, 50 mM KCH3COO, 2 mM Mg(CH3COO)_2_, 1 mM EGTA, and 10% glycerol, pH = 7.6). The focal position was identified and secured in place with an autofocus system based on total internal reflection for the entire measurement period. All data were collected at an acquisition rate of 1 kHz for 100 s using AcquireMP (Refeyn) and subsequently analyzed with DiscoverMP (Refeyn).

### Microtubule preparation

Porcine brain tubulin was isolated using the high-molarity PIPES procedure and then labeled with biotin NHS ester, Dylight-405 NHS ester, or Alexa647 NHS ester, as described previously (https://mitchison.hms.harvard.edu/files/mitchisonlab/files/labeling_tubulin_and_quantifying_labeling_stoichiometry.pdf). Pig brains were obtained from a local abattoir and used within ∼4 hrs. after death. To polymerize microtubules, 50 mM of unlabeled tubulin, 10 μM of biotin-labeled tubulin and 3.5 μM of Dylight-405-labeled (or Alexa647-labeled) tubulin were incubated with 2 mM of GTP for 20 min at 37°C. Polymerized microtubules were stabilized by the addition of 20 μM taxol (Fisher Scientific) and incubated an additional 20 min at room temperature. Microtubules were pelleted at 20, 000 x g by centrifugation through a 150 μl of 25% sucrose cushion and the pellet was resuspended in 50 μl BRB80 (80 mM Pipes (pH 6.8), 1 mM MgCl_2_ and 1 mM EGTA) containing 10 μM taxol.

### Actin preparation

Actin was purified from rabbit skeletal muscle acetone powder (Pel-Freez, Rogers, AR), as previously described (*73*). Briefly, actin was extracted in G-buffer (3 mM Tris-HCl, 0.1 mM CaCl2, 0.2 mM DTT and 0.2 mM ATP, pH 8.0) from acetone powder and centrifuged in a Fiberlite F14 6×250 LE rotor (ThermoFisher Scientific) at 10,000 rpm for 30 min at 4 ºC. The actin was polymerized by adding KCl and MgCl_2_ to final concentration at 50 mM and 2 mM, respectively. Following polymerization, KCl concentration was increased to 0.6 M and centrifuged in Beckman 45 Ti rotor at 45,000 rpm for 2 hours at 4 ºC. Pelleted actin was dialyzed against G-buffer for two days, spun in Beckman MLA50 rotor at 50,000 rpm for 20 min at 4ºC, and the resulting supernatant was snap frozen and stored at -80ºC.For labeling actin, actin was dialyzed against G-buffer pH 7.5, substituting Tris and DTT with HEPES and TCEP, respectively. Dialyzed actin was polymerized by adding KCl and MgCl_2_ to a final concentration of 50 mM and 2 mM, respectively, EZ-Link NHS-PEG4-biotin or DyLight 650 NHS ester (both from ThermoFisher Scientific) was also added to the solution. Labeled actin was sedimented at 80,000 rpm for 20 min in a Beckman TLA100.4 rotor (Beckman Coulter, Brea, CA), and dialyzed against G-buffer. Dialyzed actin was then centrifuged, aliquoted, snap frozen in liquid nitrogen and stored in -80ºC.

### Low-speed Co-sedimentation assay

Actin filaments were polymerized from 10 μM actin monomers in KMEI buffer (10 mM Imidazole, 50 mM KCl, 1 mM EGTA, and 1 mM MgCl_2_, pH8) and stabilized with 10 μM phalloidin (Fisher Scientific). Next, 5 μM F-actin was incubated with or without 1 μM mN2G^SR^ or mN2G^CH^ in SRP90 buffer for 1 hr at room temperature. The protein mixture was then centrifuged at 10,000 x g for 30 min at 4 ºC, and the resulting supernatant and pellet were examined by SDS-PAGE.

### TIRF Assays

TIRF chambers were assembled by combining an acid-washed glass coverslip prepared as previously described (http://labs.bio.unc.edu/Salmon/protocolscoverslip-preps.html), a pre-cleaned slide (Fisher Scientific), and double-sided sticky tape. The chambers were first incubated with 0.5 mg/ml PLL-PEG-biotin (Surface Solutions Group. Chicago, IL) for 5-10 min, followed by 0.5 mg/ml streptavidin (ThermoFisher Scientific for 5 min at room temperature.

Microtubules were diluted in BRB80 containing 10 μM taxol. To adhere diluted microtubules to the glass coverslip in a TIRF chamber, they were were flowed into streptavidin-adsorbed flow chambers and incubated for 2-5 min at room temperature for adhesion. To remove unbound microtubules from the chamber, the chambers were subsequently washed with SRP90 assay buffer (90 mM HEPES, 50 mM KCH3COO, 2 mM Mg(CH3COO)2, 1 mM EGTA, 10% glycerol, pH = 7.6), supplemented with 1 mg/mL BSA, 0.05 mg/mL biotin–BSA, 0.2 mg/mL K-casein, 0.5% Pluronic F-127, 10 μM taxol, and 1 μM Phalloidin pH = 7.0). Purified motor proteins were diluted to indicated concentrations in the assay buffer with 2 mM of corresponding nucleotides (ATP-Chem Imex, ADP-Sigma Aldrich, or AMPPNP-Roche Diagnostics GmbH)fr, then, the solution was flowed into the TIRF chamber.

In the KMEI buffer, 10 μM actin consisting of 30% DyLight-650-labeled and optionally 5% biotinylated actin, was polymerized and stabilized with 10 μM phalloidin, unless stated otherwise. Actin filaments were diluted in SRP90 buffer containing 1 μM Phalloidin just prior to the experiments. To anchor actin filaments to the glass surface of the TIRF chamber, streptavidin-coated chambers were first incubated with the blocking buffer (SRP90 buffer supplemented with 0.2 mg/ml K-casein, and 0.5% Pluronic F-127) for 5 mins. Then, diluted actin filaments were flowed into the TIRF chambers and incubated for 1 min at room temperature for their adhesion to the glass coverslip. Unbound actin was subsequently washed out with SRP90 assay buffer. Purified mN2G proteins were diluted as indicated in the assay buffer and subsequently flowed into the TIRF chamber.

For visualizing mN2G-mediated actin-bundling, 5 μM of F-actin (30% DyLight-650-labeled, without biotinylated actin) was incubated with 1 μM mN2G^SR^ or mN2G^CH^ in SRP90 buffer at room temperature for 10 min and then diluted 40 times in SRP90 buffer.

For actin bundle co-assembly experiments, 50 nM of phalloidin-stabilized F-actin was mixed with 25 nM mN2G^SR^ and 25 nM TMR-labeled Fascin, or 50 nM mN2G^SR^ and 50 nM TMR-labeled α-actinin, or 25 nM α-actinin-sfGFP and 25 nM TMR-labeled Fascin in SRP90 assay buffer supplemented with 0.5% methylcellulose 1500 cp and flushed into TIRF chamber. Images were acquired after >5 min incubation at room temperature.

All the images were acquired using a NIS Elements software (Nikon, Melville, NY)-controlled Nikon TE microscope using a 1.49 numerical aperture PlanApo 100× objective equipped with a TIRF illuminator (LU-N4) and an Andor iXon CCD EM camera (Oxford Instruments, Abingdon, Oxfordshire, UK). Data were analyzed manually using ImageJ (Fiji).

### Statistical analysis

The figure legends comprehensively detail all quantification and statistical analyses conducted in this study, specifying the statistical tests employed and providing exact values of “N” or “n” used in this work. Here, “N” denotes the number of trials, while “n” represents the number of data points (e.g, molecules or filaments) analyzed. Fluorescence intensities measured on the filaments were background-subtracted for each filament and assessed using either a two-tailed unpaired Student’s t-test (for comparisons between groups) or a one-way ANOVA (for comparisons involving more than two groups). Statistical analyses were performed using Prism 9 (GraphPad, San Diego, CA). Significance levels are stated on the figure. For Figure 3 Fold assays the negative control fold was set to 1.

### Alphafold modeling

The AlphaFold2 models of mN2G^SR^ and mN2G^CH^ were generated using ColabFold version 1.5.2 (Mirdita et al. 2022) with the “alphafold2_ptm” model with no templates. Five models were generated per construct, and the top-scoring model (i.e., the highest pLDDT value) was selected and relaxed. To address disordered regions in proteins, dihedral angles (phi=-165°, psi=165°) were assigned with a slight amount of Gaussian noise (+/-3°) using a custom Python script in ChimeraX (Meng et al. 2023) available at <https://github.com/delnatan/colabfold_chunker_utils/blob/cc0fc38323fae7afbabb6a3691175e00d78b82da/straighten.py>. Random noise was added to generate variations of the disordered region with each execution of the command. Visual inspection was conducted and one variant was chosen for inclusion in the final figure.

**Figure S1:**
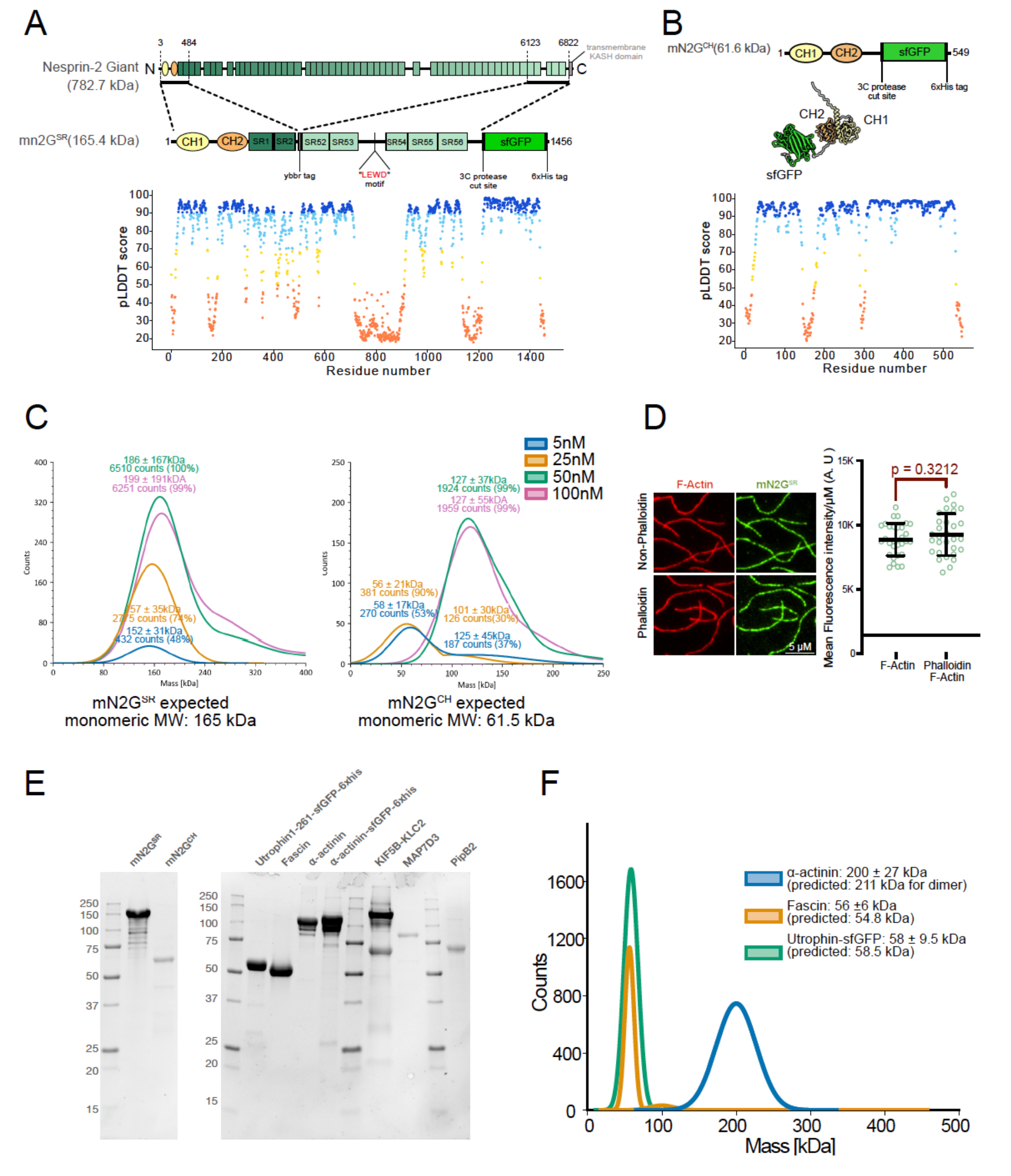
Characterization of Proteins Used in this Study. (A) Schematic domain diagrams showing parts of N2G used to construct mN2G^SR^ used in this study. The domain boundaries are drawn according to annotations in the UniProt database. A plot of pLDDT scores per-residue from AlphaFold2 is shown below for the predicted structure of mN2G^SR^ (a.a. 1-1456). (B) Schematic domain diagrams for the mN2G^CH^ construct used in this study and the cartoon rendering of its predicted structure from AlphaFold2. A plot of the pLDDT scores per-residue is shown below (a.a. 1-549). (C) Gaussian fits of the histograms of measured molecular weight of indicated proteins in solution using mass photometry at different concentrations. The calculated mass and percentage of particles within the peaks are indicated. For mN2G^SR^, n = 1231, 4354, 7256, and 7247 molecules (at 5,25,50 and 100nM, respectively) and for mN2G^CH^, n = 852, 559, 3470, and 3198 molecules (at 5, 25, 50, and 100nM respectively). (D) Mean fluorescence intensity measurements of bound 50nM mN2G^SR^ on phalloidin vs. non-phalloidin stabilized actin filaments (N=3, n=30 actin filaments for each concentration). Representative TIRF images of 50nM mN2G^SR^ on each type of actin filament. (E) Coomassie Brilliant Blue stained gel showing the purified proteins used in this study. (F) Mass photometry analysis confirming both utrophin 1-261 and fascin are monomeric, and α-actinin is dimeric. Gaussian fit of each species and derived molecular mass along with predicted mass are shown. n=6626, 3018, and 8594 for utrophin, fascin, and α-actinin, respectively.

**Figure S2:**
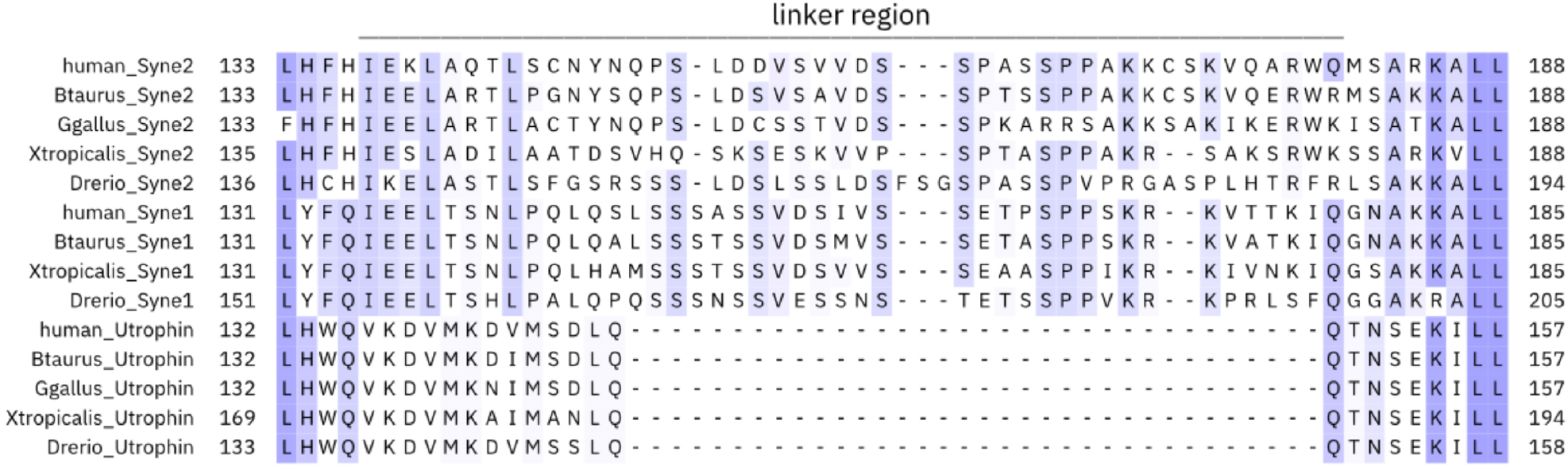
Comparison of linker regions between the tandem CH domains of nesprin-1 and -2 compared to utrophin. Multiple sequence alignment of nesprin-1, nesprin-2 and utrophin across various species: human, cow (*Bos taurus*), chicken (*Gallus gallus*), frog (*Xenopus tropicalis*), and fish (*Danio rerio*). Nesprin-1 from chicken is omitted because no sequence containing a tandem CH domain could be found in UniProt. The linker region is highlighted by a horizontal line above the alignment. Each sequence’s native numbering is shown on the left (N-terminus) and right (C-terminus) of the alignment. Nesprin-1 and nesprin-2’s linker regions have some compositional conservations and are longer than those found in utrophin (*52*).

**Figure S3:**
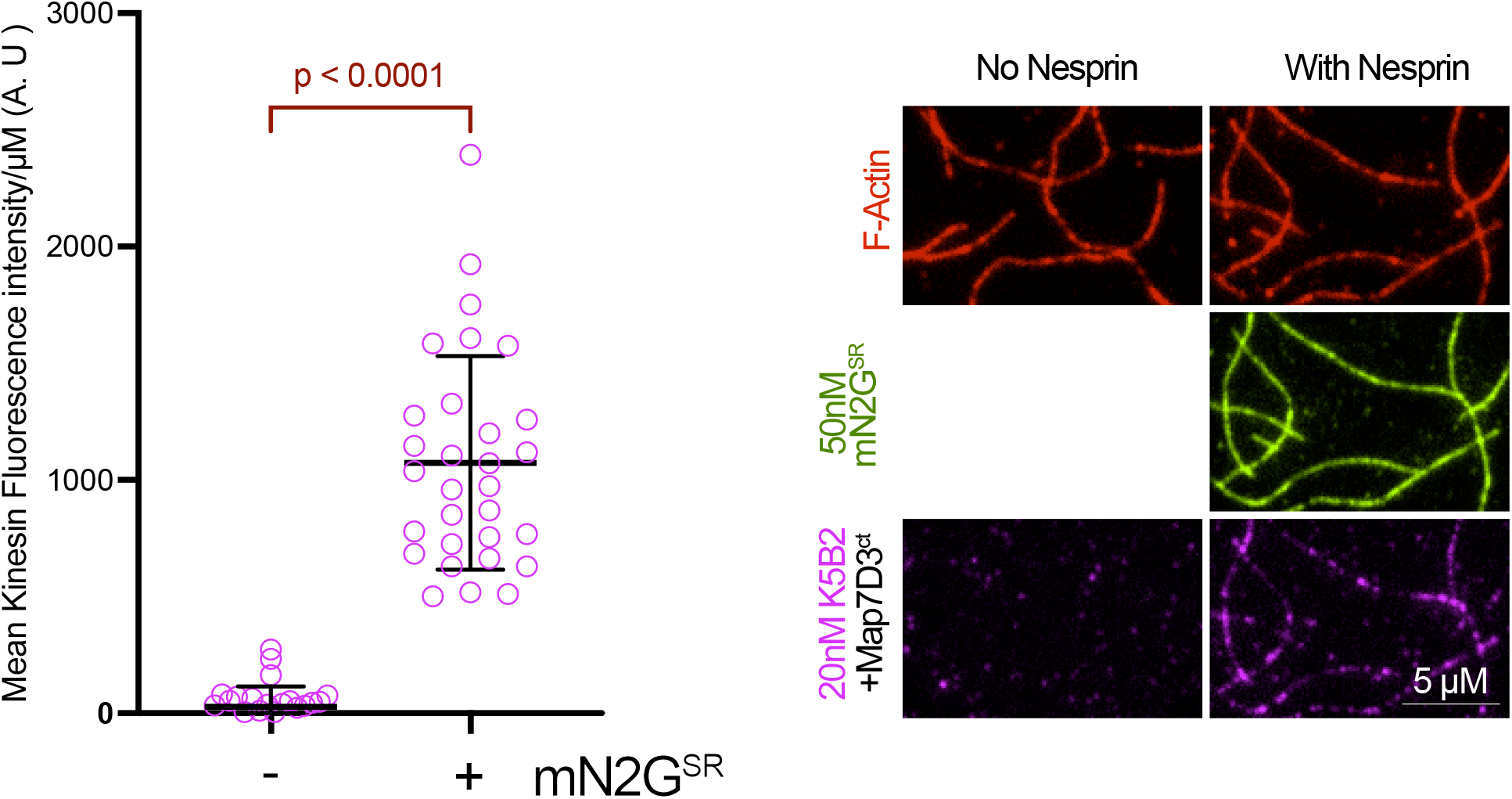
The Kinesin-Map7D3^CT^ complex does not interact with Actin without the Nesprin. Representative TIRF images and mean fluorescence intensity measurements of 20nM K5B2 (mScarlet)-50nM MAP7D3^ct^ (dark) complex with or without mN2G^SR^ binding to actin filaments (N=1, n=30 actin filaments for each concentration).

## Supplemental Movie Legends

**Movie S1: The Kinesin-mN2G**^**SR**^**-MAP7D3**^**CT**^ **Complex Recruits F-actin to Microtubules**. Representative time-lapse shows the dynamics of F-actin filaments (red) in the presence of surface-immobilized microtubules (blue), kinesin heterotetramer (magenta) and mN2G^SR^ (green), with or without the addition of MAP7D3^CT^ (unlabeled). Scale bar and time are noted in the movie.

**Movie S2: Examples of Various F-actin Behaviors During Transport Along Microtubules**. Representative time-lapse shows transport of F-actin filaments (red) along surface-immobilized microtubules microtubules (blue) in the presence of kinesin heterotetramer (magenta) and mN2G^SR^ (green). The movies show examples of various commonly observed behaviors during F-actin transport including the merging of F-acitn filaments, splitting of filaments, and filaments switching between different microtubule tracks. Scale bar and time are noted in the movie.

**Movie S3: Examples of F-actin Accumulation at Microtubule Ends and Its Effects**. Representative time-lapse shows transport of F-actin filaments (red) along surface-immobilized microtubules (blue) in the presence of kinesin heterotetramer (magenta) and mN2G^SR^ (green). The movies show examples of common accumulations of F-actin filaments at microtubule ends after processive transport. Occasionally, when F-actin accumulations are near another microtubule, they act as a bridging structure between both microtubules that can exert force and drag the end of one microtubule along the other. Surface immobilization of the microtubules prevents complete reorientation of the microtubules. Scale bar and time are noted in the movie. Scale bar and time are noted in the movie.

## Notes

### Competing Interest Statement

The authors have declared no competing interest.

